# Generalist vs specialist strategy shapes microbiomes in blood feeding parasite *Polyplax serrata*

**DOI:** 10.1101/2025.09.25.678468

**Authors:** Damián Dedecius, Jakub Kolář, Jana Martinů, Jan Štefka, Eva Nováková, Václav Hypša

## Abstract

Insects live in association with bacterial communities, collectively referred to as the microbiome. Microbiome composition varies widely across insect taxa and is shaped by multiple factors, including host phylogeny, environmental conditions, geographic distribution, and nutritional ecology. One hypothesis is that microbiome composition may also reflect whether the host adopts a generalist or specialist ecological strategy. We tested this hypothesis using the sucking louse *Polyplax serrata*, which offers several advantages as a model system. First, as permanent ectoparasites, lice inhabit a relatively stable and simplified environment, thereby minimizing potential confounding variables. Second, within *P. serrata*, two closely related lineages have been identified: one restricted to a single rodent host (*Apodemus flavicollis*), and the other exploiting two hosts (*A. flavicollis* and *A. sylvaticus*). We analyzed and compared microbiome structure in these two lineages using 16S rRNA gene amplicon sequencing. While alpha diversity did not differ between the lineages, beta diversity differed significantly, particularly in pairwise dissimilarities among individual samples. These results suggest that in *P. serrata*, host specialization strategy influences microbiome diversity, with the “generalist” lineage harboring more heterogeneous communities. This finding extends previous observations on ecological divergence between the two lineages, showing that closely related cryptic species with highly similar genomes, living sympatrically in the same environment, can rapidly evolve distinct life strategies that, in turn, shape both their genetic structure and their microbiomes.

## Introduction

Insects live in close association with bacteria acquired from their environment and diet. These microbial partners perform a range of functions that are essential for host survival and fitness (Gupta and Nair 2020). The composition of these bacterial communities, collectively referred to as the microbiome, varies widely across insect taxa (Lange et al. 2023). Microbiome structure is shaped by multiple factors, including the host’s phylogenetic relationships, environmental conditions, geography, and nutritional requirements (Jackson et al. 2023, Martoni et al. 2023, Serrato-Salas and Gendrin 2023, Yun et al. 2014). An interesting hypothesis emerging from several studies is that microbiome composition may also reflect whether the host follows a generalist or specialist ecological strategy (Brunetti et al. 2022). However, disentangling the effects of these various determinants remains challenging due to the high variability of environmental conditions.

Permanent ectoparasites that live exclusively on their hosts and feed on blood throughout their entire life cycle, such as sucking lice (Anoplura), offer a considerably simplified system for studying host–microbe interactions. Their intimate and highly specialized relationship with vertebrate host, along with strict host specificity, suggests that the host-driven factors play a major role in shaping the louse microbiome. This idea is supported by recent findings of similar bacteria in rodent fur and their lice (Ríhová et al. 2024). As in other insects that feed exclusively on vertebrate blood, the microbiome of lice is dominated by an obligate mutualistic bacterium that supplies essential nutrients missing from the blood diet, particularly B vitamins (Duron and Gottlieb 2020). Consequently, most studies to date have focused primarily on these obligate nutritional symbionts (Aksoy 1995, Sasaki-Fukatsu et al. 2006, Rihova et al. 2017, Rihova et al. 2021, Rihova et al. 2022, Rihova, Vodicka and Hypsa 2025, Martin Ríhová et al. 2023). Only recently has the full microbiome diversity been explored in a few louse species, and even then, not in relation to the host range and spectrum (Dona et al. 2021, Deng et al. 2024).

The sucking louse *Polyplax serrata* provides an excellent model for studying the relationships between microbiome structure and its potential determinants. Although formally classified as a single species, *P. serrata* comprises a complex assemblage of lineages with distinct distributions and ecological characteristics (Stefka and Hypsa 2008, Martinu et al. 2020, Martinu, Hypsa and Stefka 2018, Martinu et al. 2025). Phylogenetic analyses suggest that this assemblage includes at least two closely related but genetically distinct species that occur across much of Europe. In both previous research and the present study, these are referred to as N (nonspecific) and S (specific) lineages. The primary distinction between them lies in host specificity: the S lineage is a strict specialist restricted to *Apodemus flavicollis*, while the more generalist N lineage can also parasitize *A. sylvaticus*. Although sucking lice are generally known for narrow host specificity, they are not always confined to a single host species, and the broader host range of the N lineage is thus not unprecedented. Many louse species are oligoxenous, parasitizing several (often closely related) host species (Durden and Musser 1994). Moreover, host range can expand rapidly if a louse species is introduced into a new region (Wang, Durden and Shao 2020). In addition to the major division between the N and S lineages, further subdivision is evident within the S lineage. Unlike the panmictic structure of the N lineage, the S lineage is divided into two geographically distinct sublineages: SW (S West) and SE (S East), which are separated by a narrow hybrid zone.

Previous comparative studies of the two major lineages have shown that the genomes of the N and S lineages are highly similar, with complete synteny interrupted only by a single small translocation (Martinu et al. 2023). In contrast, ecological differences specifically the specialist versus generalist strategies) are reflected in their different prevalences on the shared host *Apodemus flavicollis* (Martinu et al. 2025). In the present study, we build on these findings by examining the microbiome compositions of these two ecologically distinct but genetically closely related lineages. Our primary hypothesis is that the difference in host-use strategy plays a key role in shaping their microbiomes. However, we also consider and discuss additional factors that may influence microbiome structure.

## Results and discussion

### EMicrobiome richness and composition

The amplicon screening identified average of 68 distinct OTUs per sample, showing that despite their limited access to diverse environments, the permanent blood-feeding ectoparasites *P. serrata* harbor a relatively rich microbiome (Figure 1A, Supplementary table S1). Placing this number in the context of other exclusive blood feeders is difficult due to scarcity of comparable studies. Moreover, evaluating microbiome richness in these groups is further complicated by the dominance of one or few symbionts. For example, in tsetse flies, 99,7% of amplicon reads were assigned to the three known symbionts *Wigglesworthia, Sodalis*, and *Wolbachia* (Gaithuma et al. 2020). Similarly, in the bedbug *Cimex hemipterus*, the two most abundant symbionts, *Wolbachia* and *Symbiopectobacterium*, accounted for up to 99% of the reads (Lim and Ab Majid 2021). In our data, the most abundant OTU1 was assigned to the genus *Legionella*, clearly corresponding to the obligate symbiont *L. polyplacis* (Rihova et al. 2017). It was detected in the majority of the samples; however, its read abundance varied substantially, ranging from nearly 100% to near or complete absence in few samples (Figure 1; Supplementary table S1).

**Figure 1.**
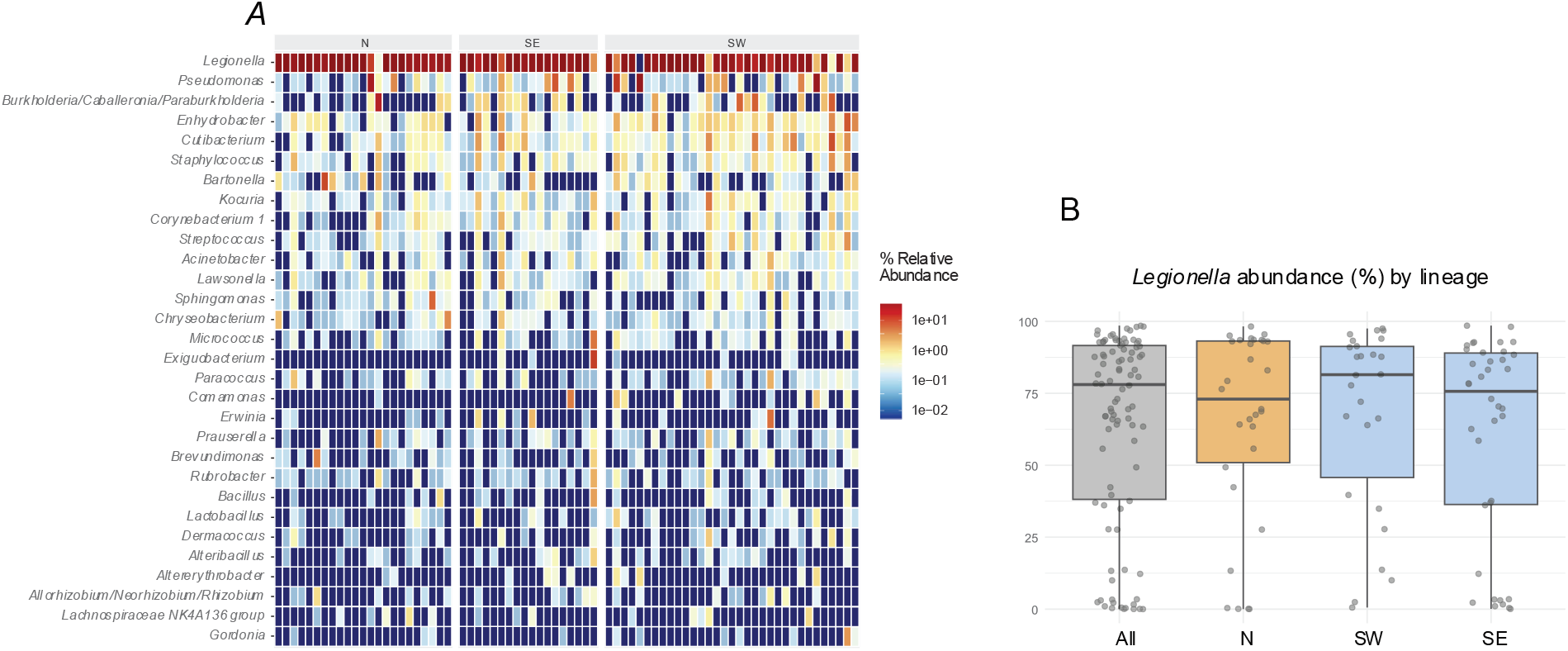
Microbiome composition based on 16S rRNA gene amplicon data. (A). Relative abundance of the 30 most abundant taxa. (B) Distribution of Legionella abundance (read counts) across samples. Boxplots show distributions for all samples, the N lineage, and the S lineage, the latter divided into two sublineages (SE = S east, SW = S west; see legend of Figure 2 for details).

Such variation in the proportion of reads assigned to an obligate symbiont across individual hosts (with some samples even lacking detectable symbiont reads) has also been reported in human lice *Pediculus humanus* (Agany et al. 2020). Because obligate symbionts are essential for their hosts, these differences likely reflect the physiological or developmental stage of the host, rather than factors driving microbiome variation examined in our study. This interpretation is supported by the read distribution across our samples (Figure 1B), which is continuous between 0 and 100%, rather than binary (present/absent). As explained in Methods, OTU1 was excluded from subsequent analyses to avoid introducing of misleading signal.

### Microbiome diversities

The global PERMANOVA model, incorporating Lineage, Host, and Region as explanatory variables (see Figure 2 for the complete statistical workflow), explained approximately 14% of the total variation in microbiome structure (p = 0.008, R^2^ = 0.14, F = 1.26). This indicates a moderate but statistically significant combined effect of the three predictors. Marginal tests identified Lineage (N versus S) as the primary driver of variation (p = 0.016), while Region (p = 0.046) showed a borderline significant effect, and Host (p = 0.178) did not reach statistical significance. Consequently, follow-up analyses focused on comparing diversity measures between the N and S lineage microbiomes. These analyses revealed that the two lineages exhibit similar alpha diversity, as measured by the Shannon index (p = 0.70; Figure 3A), indicating no significant difference in overall richness and evenness of their microbiomes. In contrast, microbiome composition differed significantly between lineages (PERMANOVA, Bray–Curtis, p = 0.016, R^2^ = 0.06, F = 1.96; Figure 3B). This difference may be influenced by greater within-lineage variability in the N lineage (Wilcoxon test on pairwise dissimilarities, p = 6.01 x 1^-8^; Figure 3C). However, a formal test of multivariate dispersion (PERMDISP) did not reach significance (p = 0.089; Figure 3D), providing only marginal evidence for differences in within-group variability. Overall, these results suggest that the S lineage tends to harbor more homogeneous microbiomes, whereas the N lineage may exhibit greater heterogeneity.

**Figure 2.**
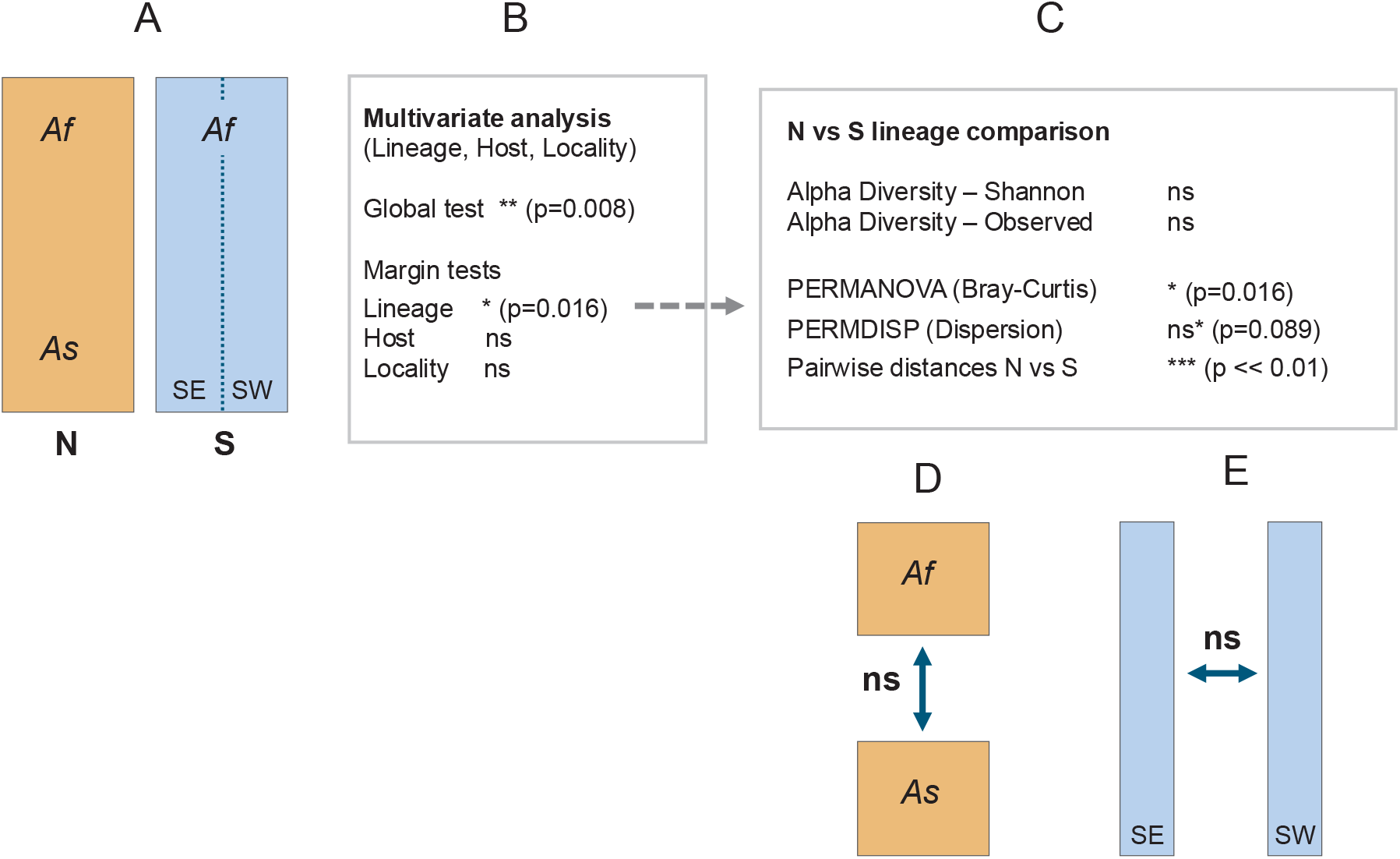
Schematic overview of statistical analyses and their outcomes across the N (grey) and S (blue) lineages, from the two hosts: Apodemus flavicollis (Af) and A. sylvaticus (As). Signiicance levels are indicated as * (p < 0.05), ** (P < 0.001), and ns (p > 0.05); ns* marks a nonsigniicant but borderline result for beta diversity distribution. For the signiicant results discussed in the main text, exact values are provided next to the corresponding symbols.

**Figure 3.**
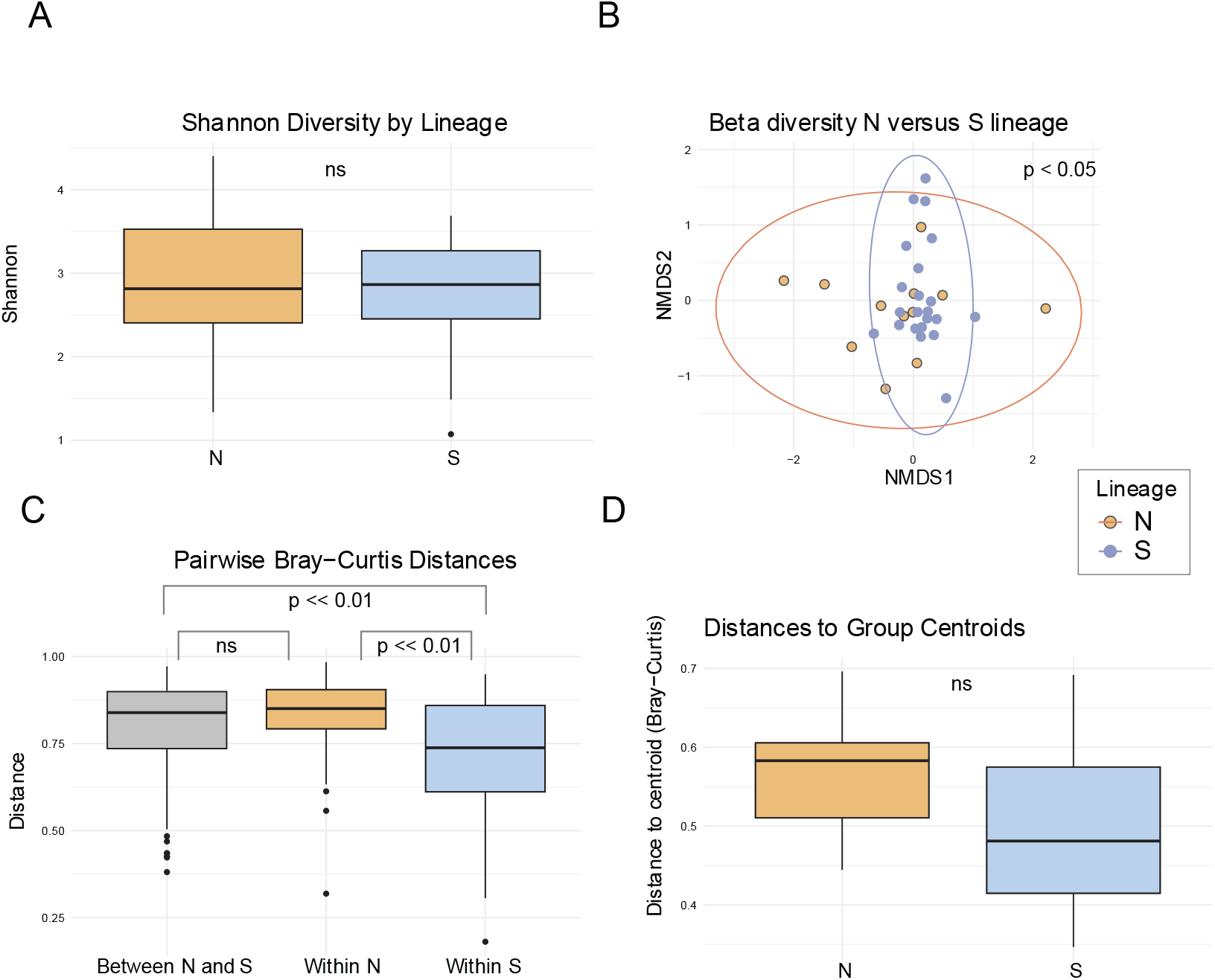
Comparison of microbiome diversity between N and S lineages. (A) Boxplots of alpha diversity (Shannon index). (B) Scatter plot of beta diversity with 95% conidence ellipses, analyzed by PERMANOVA based on Bray–Curtis distances (p = 0.016). (C) Boxplots of beta diversity based on pairwise Bray–Curtis distances (p = 6.01 × 10‐8). (D) Boxplots of beta diversity dispersion analyzed using PERMDISP (p = 0.089).

As discussed above, investigating microbiomes in obligate blood-feeders is complicated by the dominance of obligate nutritional symbionts. In our study, all analyses were performed after excluding *L. polyplacis* (see also Materials and Methods), which resulted in a substantial reduction in data. This necessitates careful consideration of statistical power in subsequent analyses. The key finding that the N and S lineages differ in microbiome beta diversity is based on the test with statistical power ≈ 0.39, which is considerably below the commonly recommended threshold of 0.80. Nevertheless, several lines of evidence support the validity of the observed pattern. First, low statistical power is primarily a concern because it increases the risk of failing to detect true effects (Type II error). However, in this case, we are evaluating the reliability of an effect that was detected. This finding is supported by a statistically significant p-value of PERMANOVA analysis (0.016), and even more strongly by the extremely low p-value observed for pairwise distances within N versus S lineages (p = 6.01 × 10^-8^). Therefore, despite the low power, the robustness of the observed signal (evidenced by the strength of the p-values) suggests that the detected difference in microbiome beta diversity between the lineages is unlikely to be a false positive. Second, when calculated using Jaccard rather than Bray-Curtis distances, PERMANOVA analysis yielded significant result with an even lower p-value (0.009). Finally, an auxiliary analysis using non-rarefied data (which retained a larger number of samples) yielded similar results (see Supplementary table S2). This further supports the consistency of the main signal, even though the non-rarefied dataset may be subject to additional sources of bias or artifacts.

Numerous factors can influence differences in microbiome structure across species or populations, including geography, trophic status, dietary source, and environment. Among these, geography was explicitly tested in our multivariate model and showed borderline statistical significance (p = 0.046). Its effect was notably weaker compared to that of Lineage. This corresponds well with the geographical structure of our sampling. Most samples come from several localities in Bohemia, all of the localities contain both lineages. There are two exceptions: the populations from Bavaria and Saxony contain only N lineage lice. However, when the Bavaria and Saxony samples were removed from the analysis, the difference between the N and S remained significant (Supplementary figure S1).

Another potential factor influencing microbiome composition is the trophic status of the insect host. In the blood-feeding species *Cimex hemipterus*, it has been shown that microbiome composition varies with trophic state: starved individuals possess a richer microbiome than blood-fed ones. However, trophic status can very likely be excluded as an explanation for our results. A crucial difference exists between the feeding strategies of bed bugs and sucking lice. Bed bugs feed only once every few days (Reinhardt and Siva-Jothy 2007), experiencing a cyclical shift between fed and starved states. In contrast, sucking lice feed multiple times per day (Schaub, Kollien and Balczun 2012), resulting in a more stable trophic state. More importantly, since the only significant predictor of microbiome variation identified in our analysis was the division between the N and S lineages, it is unlikely that trophic status would follow this same pattern. As explained above, trophic status is likely reflected in the abundance of the obligate symbiont *L. polyplacis*, which was excluded from the analysis for this very reason.

The most likely explanation for the observed microbiome differences is ecological variation, specifically host specificity. The S lineage is restricted to a single rodent host, *Apodemus flavicollis*, while the N lineage parasitizes two rodent species, *A. flavicollis* and *A. sylvaticus*. Mapping rodent hosts on genetic structure of the N lineage confirms that these lice frequently switch from one host species to another, rather than forming host specific clusters (see details below; Figure 4). This difference in host usage may influence microbiome composition in two ways. First, microbiome composition may reflect the *current host*, meaning it can change rapidly, in each louse individual being determined by the host on which the louse is presently residing. Second, in a longer ecological time, host switching creates a more dynamic environment, leading to more diverse microbiomes. This factor we term as the *host-usage strategy*. A targeted comparison of N lineage lice from the two host species revealed no significant difference in microbiome composition. Samples from *A. flavicollis* and *A. sylvaticus* were intermixed in ordination plots, and statistical analysis yielded non-significant results (p = 0.28; Supplementary figure S1). These findings do not support the *current host* hypothesis.

**Figure 4.**
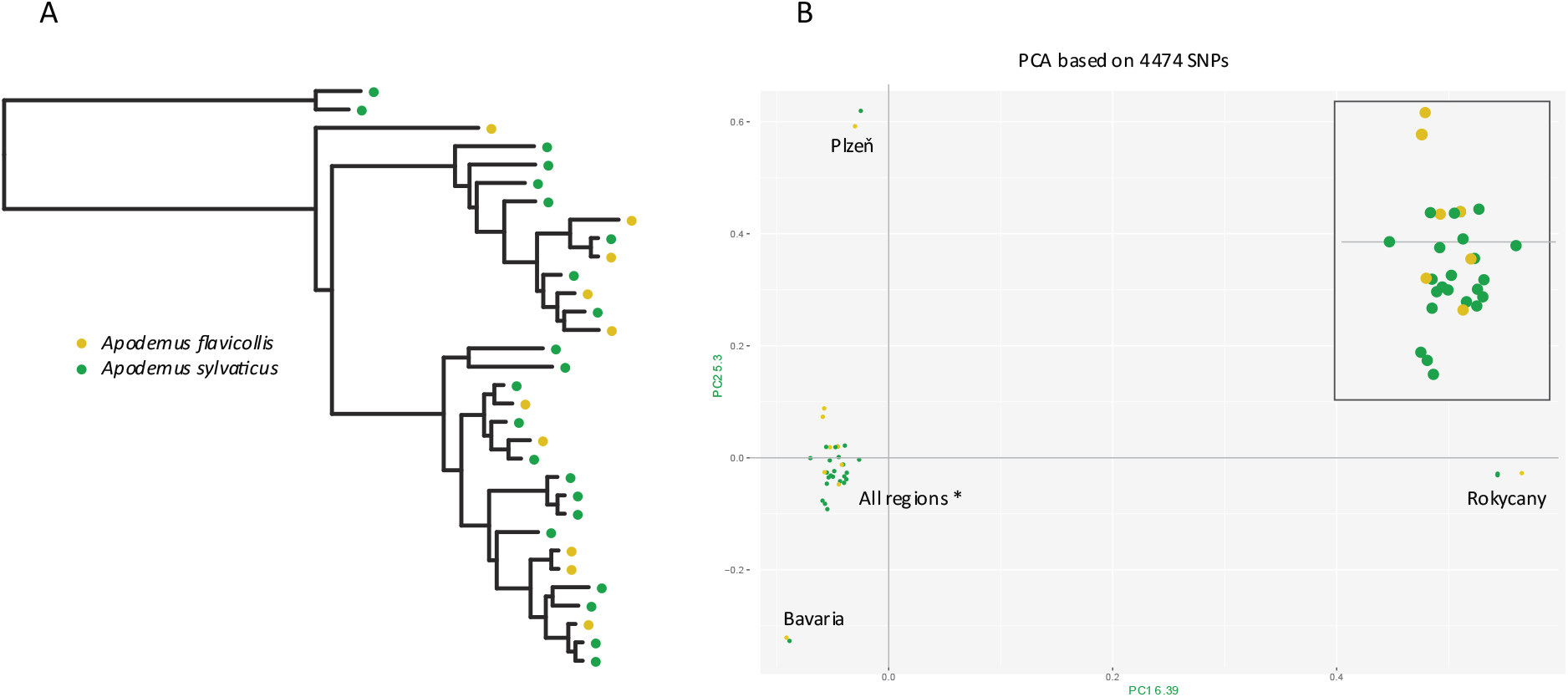
Rodent host distribution mapped onto the genetic structure of the N lineage. (A) Maximum-likelihood phylogenetic tree inferred with IQ-TREE from a concatenated alignment of eight mitochondrial minichromosomes (10,754 positions; partitions and substitution models listed in Supplementary Data). (B) Principal Component Analysis (PCA) based on SNP data. Both host species are intermixed across all four genetic clusters (Plzen, Rokycany, Bavaria, and a large cluster labeled “All regions*” that includes individuals from all sampled sites). For clarity, a zoomed-in view of the large cluster is shown in the upper right corner.

The *host-usage strategy*, specifically, the broader spectrum of rodent hosts exploited by the N lineage, offers the most plausible explanation for the observed microbiome differences. From a biological perspective, frequent switching between rodent host likely exposes N lineage lice to more complex and variable environmental conditions, which in turn promotes greater microbiome diversity over time. Due to these different strategies and distinct evolutionary histories, the S and N lineages also exhibit divergent genetic structures. Based on single nucleotide polymorphism (SNP) data, the N lineage displays significantly higher nucleotide diversity than the S lineage (0.051 vs. 0.006). This finding is consistent with Nadler’s hypothesis (Nadler 1995) or a similar Specialist-Generalist Variation Hypothesis (Li et al. 2014), which posit that generalists, owing to more frequent opportunities for dispersal, tend to maintain populations with higher local diversity (Li et al. 2014). This raises the key question of whether microbiome diversity differences are driven directly by the *host-usage strategy* (as the ultimate factor shaping ecological exposure) or indirectly through the distinct genetic structures of the N and S lineages (which themselves are consequences of the *host-usage strategies*). To address this, we compared pairwise Bray-Curtis distances derived from microbiomes with Euclidean distances derived from SNPs. The Mantel test revealed very weak correlations, with borderline nonsignificance (N lineage: r=0.20, p=0.61; S lineage: r=0.07, p=0.055). Based on these results and the overall biology of the system, we propose that the direct effect of *host-usage strategies* provides the more plausible explanation for the microbiome diversity differences.

### Host independent structure of the N lineage

The interpretation provided above assumes a real “generalistic” nature of the N lineage, that is, frequent random switches between the two *Apodemus* species (as contrast to few host specific clusters). This assumption has been confirmed by both phylogeny based on mitochondrial genome and population structure derived from genome-wide SNPs (Figure 4). In the phylogenetic tree, lice from different hosts are interspersed across the topology with only occasional monophyly of samples from the same host. Similarly, in PCA, most cluster without host-specific division. A few samples were positioned far from the main cluster by large distances on both axes, but each of these small outlier groups contained samples from both host species.

### Conclusion

Our analyses suggest that in *P. serrata* lice, the specialist versus “generalist” strategy (i.e., exploiting a single versus two host species) influences microbiome diversity, with the “generalist” lineage harboring more heterogeneous microbiomes. This result fits into a broader pattern observed across insect studies, where similar ecological factors play an important role in shaping the evolution of microbiome diversity. For example, in Chrysomelidae, generalist species harbor more diverse microbiomes than specialists (Brunetti et al. 2022). Within Anoplura, significant difference was detected in human louse *Pediculus humanus* between the head and body ecotypes, even though these ecotypes do not form mutually exclusive monophyletic clusters (Agany et al. 2020). With respect to our model species *P. serrata*, the findings reported here extend the previous observations on ecological differences between the N and S lineages. Taken together, the accumulated results illustrate how closely related cryptic species with highly similar genomes, living in sympatry in the same environment, can rapidly evolve different life strategies that, in turn, shape both their genetic structure and the composition of their microbiomes.

## Material and Methods

### Samples and DNA preparation

Sucking lice *Polyplax serrata* were collected from *Apodemus* field mice captured in the northwest of the Czech Republic (CZ) and at several German localities (Supplementary table S1). Permission for field work was obtained from the Committee on the Ethics of Animal Experiments of the University of South Bohemia, the Ministry of the Environment of the Czech Republic, and by the Ministry of the Agriculture of the Czech Republic (Nos. MZP/2017/630/854, 43873/2019-MZE-18134, MZP/2021/630/2459). Mice were captured using wooden snap traps. Lice were brushed from the fur and stored in 100% ethanol at −20 °C. DNA was extracted from individual lice using the Qiagen QIAamp DNA Micro Kit (Qiagen). Lineage assignment (S or N) was determined by sequencing a 379 bp fragment of the mitochondrial cytochrome oxidase subunit I gene (COI), as described in detail in (Martinu, Hypsa and Stefka 2018).

### Amplicon sequencing and downstream processing

The V3–V4 region of the 16S rRNA gene was amplified from all samples using the QIAGEN Multiplex PCR Kit (Qiagen, Hilden, Germany). Amplifications included four blank (negative) PCR controls and two commercial genomic DNA (positive) controls (ATCC® MSA-1000™ and MSA-1001™, each consisting of the same ten bacterial species in different proportions). A two-step PCR protocol was applied with primers 341F and 805R containing staggered spacers and Illumina overhang adapters, followed by an index PCR to add sample-specific barcodes (Illumina 16S Metagenomic Sequencing Library Preparation Guide). Purified amplicons were quantified, pooled equimolarly, and sequenced on an Illumina MiSeq platform using v2 chemistry with 500 cycles. Demultiplexed raw reads were processed with USEARCH v11.0.667 (Edgar 2013), including primer removal, read merging, trimming, quality filtering, and OTU clustering. Reads were merged with zero allowed mismatches, filtered using the stringent option *-fastq_maxee 1.0*, and trimmed to 400 bp. An OTU table was generated by clustering sequences at 100% identity, followed by de novo OTU picking with USEARCH global alignment at 97% identity, including chimera removal (Edgar 2013). Taxonomic assignments were performed with BLASTn against the SILVA_132 database (https://www.arb-silva.de/fileadmin/silva_databases/release_132/Exports/SILVA_132_SSURef_tax_silva_trunc.fasta.gz). Subsequent data filtering, rarefaction, and heatmap visualization were conducted in R using the microeco v0.16.0 package (Liu et al. 2021) and ggplot2 v3.4.2 (Ito and Murphy 2013). Specifically, the OTU table was filtered to exclude archaeal, eukaryotic, mitochondrial, and chloroplast sequences.

### Microbiome diversity

To compare microbiome structures, we applied several statistical tests across different combinations of louse lineages and sublineages (Figure 2). To minimize noise introduced by the obligate nutritional symbiont *Legionella polyplacis*, OTU1 assigned to this symbiont was excluded from the analyses. The rationale is as follows: in the majority of samples, more than 50% of reads were assigned to this OTU, with some samples reaching up to 99% (Figure 1, Supplementary table S1). On the other hand, there were also samples with low read counts for this OTU, or even complete absence. However, *L. polyplacis* is a nutritionally essential, obligate symbiont that is transovarially transmitted and fixed in *P. serrata* lice (Rihova et al. 2017). Its apparent absence in some samples likely reflects the physiological condition of the host louse rather than environmental variation. Given the high and variable abundance of *L. polyplacis* across samples, often dominating the read count, its inclusion would introduce a strong, misleading signal into the analysis. Therefore, excluding OTU1 ensures a more accurate analysis of factors underlying the microbiome structure.

For all analyses, the datasets were rarefied to 1 000 reads per sample, and samples with fewer reads were excluded (a parallel auxiliary analysis was done without rarefication; see Discussion). The main comparison was conducted between the N and S lineages. All analyses were performed in the R environment (Team 2014), including data cleaning, statistical testing, and result visualization. The analysis utilized the packages microeco (Liu et al. 2021), vegan (Oksanen 2025), and ggplot2 v3.4.0. (Ito and Murphy 2013). To compare microbial diversity, we first conducted multivariate analyses to assess the combined and individual effects of the variables Lineage, Host, and Locality. Based on these results, we further analyzed the effect of Lineage on alpha and beta diversity using the following methods: Shannon index and Observed richness for alpha diversity; beta diversity dissimilarity, distribution plots, PERMANOVA (based on Bray–Curtis distances), and PERMDISP (assessing dispersion using Wilcoxon tests) for beta diversity. To control for the possible effect of the within-lineage characteristics, we further compared the N lineage samples from the two different hosts (*A. flavicollis* and *A. sylvaticus*), and S samples from the two mitochondrial lineages (SW and SE).

### Relation between population structure and host specificity

To determine whether the N lineage is entirely “generalistic” or instead forms host-determined clusters, we analyzed the population structure of the lice using two complementary approaches: mitochondrial DNA-based phylogeny and genome-wide nuclear SNP-based genetic structure. For this purpose, 38 lice from both host species (*A. sylvaticus* and *A. flavicollis*) sampled from the same localities where possible, were selected and their genomes re-sequenced. DNA concentrations were verified using a Qubit 2.0 Fluorometer (Invitrogen) with high-sensitivity reagents. Ultralow Input Libraries (Tecan) were prepared, and 150-bp paired-end sequencing was carried on one S4 Illumina Novaseq 6000 flow cell at the W. M. Keck Center (University of Illinois, Urbana, IL, USA).

For SNP analysis, genomic reads were mapped to the reference genome of *Polyplax serrata* S lineage (GCA_037055385.1) using bowtie2 (Langmead and Salzberg 2012). The reference was indexed using SAMtools (Li et al. 2009), and a sequence dictionary was created with CreateSequenceDictionary in Picard version 2.0.1 (https://broadinstitute.github.io/picard/). SAM files were converted to sorted BAM files and indexed with SAMtools. Duplicated sequences were removed using Picard 2.0.1 and mapping success was assessed with qualimap (http://qualimap.bioinfo.cipf.es/). Variant calling was performed with GATK following the “Best Practices” workflow (Van der Auwera et al. 2013), and the resulting SNP set was filtered using the thresholds QD < 2.0, FS > 60.0, MQ < 40.0, and MQRankSum < −12.5. In the R package SNPRelate (https://doi.org/10.18129/B9.bioc.SNPRelate), SNPs with a minor allele frequency (MAF) < 0.05 and linkage disequilibrium (LD) > 0.2 were removed. Population structure was reconstructed using PCA within the same package.

Like in other *Polyplax* lice, the mitochondrial DNA of *P. serrata* is organized into 11 separate minichromosomes (Martinu et al. 2020). To recover these minichromosomes from N lineage lice, we performed metagenomic assemblies using SPAdes (Bankevich et al. 2012). For each assembly, we created a database using the algorithm implemented in Genious (Kearse et al. 2012). Within each database, contigs corresponding to minichromosomes were identified via BLAST, using previously published minichromosomes from the S lineage of *P. serrata* as queries (Martinu et al. 2020). The extracted minichromosomes were aligned using a codon-based algorithm implemented in Geneious, and the resulting alignments were concatenated into a single nucleotide matrix. PhyIogenetic reconstruction was performed in IQ-TREE (Trifinopoulos et al. 2016), treating each minichromosome as partitions for which IQ-TREE was used to determine the best-fitting model for each partition (Supplementary Data).

### Comparison of genetic diversities of the lice and the microbiomes

To evaluate possible effect of louse genetic diversification on microbiome beta diversity, we performed two complementary tests. First, nucleotide diversity (π) was calculated and compared between the two louse lineages. For this purpose, we used SNP datasets derived from specimens with matching microbiome profiles. For the N lineage, we selected 28 of the specimens that had been re-sequenced as described above. For S lineage, 64 specimens were sequenced on the same Illumina NovaSeq lane, and SNP datasets were generated using the same pipeline. Nucleotide diversity was then calculated separately for each lineage from the VCF file using the vcfR package in R (Knaus and Grunwald 2017). Second we assessed the correlation between louse genetic diversification and microbiome beta diversity. Raw genotype data in VCF format were first converted into PLINK binary files (.bed,.bim,.fam), and genotypes were exported into a numeric matrix using PLINK’s --recode A option (Purcell et al. 2007). Pairwise genetic distances between individuals were calculated in Python (using the Euclidean distance metric) with the pdist and squareform functions from the SciPy library (Virtanen et al. 2020). The resulting genetic matrices for the N and S lineages were compared to the Bray-Curtis distance matrices of their microbiomes using a Mantel test implemented in R with vegan package. Prior to testing, both matrices were reordered to ensure that samples were matched. The Mantel test was performed with 999 permutations using Pearson correlation to assess the relationship between genetic and microbiome dissimilarities. To visualize this relationship, the lower-triangular elements of both matrices were extracted and plotted against each other using ggplot2, with a regression line overlay and the Mantel correlation coefficient (r) and associated p-value displayed in the plot title.

## Supplements

**Supplementary figure S1**. (A) PERMANOVA analysis showing group separation after removal of Bavarian and Saxony samples. (B) PERMANOVA results comparing samples from two host species.

**Supplementary table S1:** Data used in statistical analyses. **OTU** - list of OTUs and number of reads across the samples (OTU1, assigned to *Legionella*, was removed prior to the analyses; see Material and Methods and Discussion). **Metadata** - metadata specifyng individual samples. **Taxonomy** - taxonomy list used to assign taxa to OTUs.

**Supplementary table S2:** Results of the analyses with non-rarefied dataset

**Supplementary table S3:** List of samples used in the following genetic and phylogenetic analyses.

**Supplementary data:** Concatenated matrix of minichromosomes used in the phylogenetic analysis.

## Acknowledgements

We would like to thank the students and colleagues at the Department of Parasitology, Faculty of Science, University of South Bohemia, Czech Republic, for their assistance during field sampling. Access to computing and storage facilities owned by parties and projects contributing to the National Grid Infrastructure MetaCentrum provided under the program “Projects of Large Research, Development, and Innovations Infrastructures” (CESNET LM2015042) is greatly appreciated. This work was supported by the Grant Agency of the Czech Republic (grant 21-02532S to VH).

## References

Agany, D., R. Potts, J. Hernandez, E. Gnimpieba & J. Pietri (2020) Microbiome Diferences Between Human Head and Body Lice Ecotypes Revealed by 16S RRNA Gene Amplicon Sequencing. Journal of Parasitology, 106, 14–24. doi.org/10.1645/19-132.

Aksoy, S. (1995) Wigglesworthia gen. nov. and Wigglesworthia glossinidia sp. nov., taxa consisting of the mycetocyte-associated, primary endosymbionts of tsetse flies. Int J Syst Bacteriol, 45, 848–51.

Bankevich, A., S. Nurk, D. Antipov, A. A. Gurevich, M. Dvorkin, A. S. Kulikov, V. M. Lesin, S. I. Nikolenko, S. Pham, A. D. Pr ibelski, A. V. Pyshkin, A. V. Sirotkin, N. Vyahhi, G. Tesler, M. A. Alekseyev & P. A. Pevzner (2012) SPAdes: a new genome assembly algorithm and its applications to single-cell sequencing. J Comput Biol, 19, 455–77.

Brunetti, M., G. Magoga, F. Gionechetti, A. De Biase & M. Montagna (2022) Does diet breadth afect the complexity of the phytophagous insect microbiota? The case study of Chrysomelidae. Environmental Microbiology, 24, 3565–3579.

Deng, Y., C. Yao, Y. Fu, Y. Zhuo, J. Zou, H. Pan, Y. Peng & G. Liu (2024) Analyses of the gut microbial composition of domestic pig louse Haematopinus suis. Microbial Pathogenesis, 197:107106. doi: 10.1016.doi:10.1016/.micpath.2024.107106

Dona, J., S. Herrera, T. Nyman, M. Kunnasranta & K. Johnson (2021) Patterns of Microbiome Variation Among Infrapopulations of Permanent Bloodsucking Parasites. Frontiers in Microbiology, 12, 642543. doi: 10.3389/fmicb.2021.642543.

Durden, L. & G. Musser. 1994. The sucking lice (Insecta, Anoplura) of the world: a taxonomic checklist with records of mammalian hosts and geographical distributions. In Bulletin of the AMNH in Sucking Lice and Hosts; no. 218

Duron, O. & Y. Gottlieb (2020) Convergence of Nutritional Symbioses in 0bligate Blood Feeders. Trends in Parasitology, 36, 816–825.

Gaithuma, A., J. Yamagishi, K. Hayashida, N. Kawai, B. Namangala & C. Sugimoto (2020) Blood meal sources and bacterial microbiome diversity in wild-caught tsetse flies. Scientiic Reports, 10, 5005. doi.org/10.1038/s41598-020-61817-2.

Gupta, A. & S. Nair (2020) Dynamics of Insect-Microbiome Interaction Influence Host and Microbial Symbiont. Frontiers in Microbiology, 11:1357. doi: 10.3389/fmicb.2020.01357.

Ito, K. & D. Murphy (2013) Application of ggplot2 to Pharmacometric Graphics. CPT Pharmacometrics Syst Pharmacol, 2, e79.

Jackson, R., P. Patapiou, G. Golding, H. Helantera, C. Economou, M. Chapuisat & L. Henry (2023) Evidence of phylosymbiosis in Formica ants. Frontiers in Microbiology, 4:1044286. doi: 10.3389/fmicb.2023.1044286.

Kearse, M., R. Moir, A. Wilson, S. Stones-Havas, M. Cheung, S. Sturrock, S. Buxton, A. Cooper, S. Markowitz, C. Duran, T. Thierer, B. Ashton, P. Meintes & A. Drummond (2012) Geneious Basic: An integrated and extendable desktop software platform for the organization and analysis of sequence data. Bioinformatics, 28, 1647–1649.

Knaus, B. & N. Grunwald (2017) VCFR: a package to manipulate and visualize variant call format data in R. Molecular Ecology Resources, 17, 44–53.

Lange, C., S. Boyer, T. Bezemer, M. Lefort, M. Dhami, E. Biggs, R. Groenteman, S. Fowler, Q. Paynter, A. Mogena & M. Kaltenpoth (2023) Impact of intraspecific variation in insect microbiomes on host phenotype and evolution. Isme Journal, 17, 1798–1807.

Langmead, B. & S. Salzberg (2012) Fast gapped-read alignment with Bowtie 2. Nature Methods, 9, 357–U54.

Li, H., B. Handsaker, A. Wysoker, T. Fennell, J. Ruan, N. Homer, G. Marth, G. Abecasis & R. Durbin (2009) The Sequence Alignment/Map format and SAMtools. Bioinformatics, 25, 2078–9.

Li, S., R. Jovelin, T. Yoshiga, R. Tanaka & A. Cutter (2014) Specialist versus generalist life histories and nucleotide diversity in Caenorhabditis nematodes. Proceedings of the Royal Society B-Biological Sciences, 281:20132858. doi: 10.1098/rspb.2013.2858.

Lim, L. & A. H. Ab Majid (2021) Characterization of bacterial communities associated with blood-fed and starved tropical bed bugs, Cimex hemipterus (F.) (Hemiptera): a high throughput metabarcoding analysis. Scientiic Reports, 11, 8465. doi: 10.1038/s41598-021-87946-w.

Liu, C., Y. Cui, X. Li & M. Yao (2021) microeco: an R package for data mining in microbial community ecology. Fems Microbiology Ecology. 97:fiaa255. doi: 10.1093/femsec/fiaa255.

Martin Říhová, J., S. Gupta, A. Darby, E. Nováková & V. Hypša (2023) Arsenophonus symbiosis with louse flies: multiple origins, coevolutionary dynamics, and metabolic significance. mSystems, 0, e00706–23.

Martinu, J., V. Hypsa & J. Stefka (2018) Host specificity driving genetic structure and diversity in ectoparasite populations: Coevolutionary patterns in Apodemus mice and their lice. Ecology and Evolution, 8, 10008–10022.

Martinu, J., J. Stefka, A. Poosakkannu & V. Hypsa (2020) “Parasite turnover zone” at secondary contact: A new pattern in host-parasite population genetics. Molecular Ecology, 29, 4653–4664.

Martinu, J., J. Stefka, K. Vránková & V. Hypsa (2025) Diferent life strategies of closely related louse species in sympatry: specialist and “generalist” lineages of Polyplax serrata. International Journal For Parasitology, 55, 27–34.

Martinů, J., H. Tarabai, J. Štefka & V. Hypša (2023) Highly resolved genomes as a tool for studying speciation history of two closely related louse lineages with diferent host specificities. Genome Biol Evol.16(3):evae045. doi: 10.1093/gbe/evae045.

Martoni, F., S. Bulman, A. Piper, A. Pitman, G. Taylor & K. Armstrong (2023) Insect phylogeny structures the bacterial communities in the microbiome of psyllids (Hemiptera: Psylloidea) in Aotearoa New Zealand. Plos One, 18(5):e0285587. doi: 10.1371/ournal.pone.0285587.

Nadler, S. (1995) Microevolution and the genetic structure of parasite populations. Journal of Parasitology, 81, 395–403.

Oksanen, J., Blanchet, F.G., Kindt, R., Legendre, P., Minchin, P.R., O’Hara, R.B., Simpson, G.L., Solymos, P., Stevens, M.H.H., Wagner, H. 2025. Vegan: Community ecology package. The Comprehensive R Archive Network (CRAN).

Purcell, S., B. Neale, K. Todd-Brown, L. Thomas, M. Ferreira, D. Bender, J. Maller, P. Sklar, P. de Bakker, M. Daly & P. Sham (2007) PLINK: A tool set for whole-genome association and population-based linkage analyses. American Journal of Human Genetics, 81, 559–575.

Reinhardt, K. & M. Siva-Jothy (2007) Biology of the bed bugs (Cimicidae). Annual Review of Entomology, 52, 351–374.

Rihova, J., G. Batani, S. M. Rodriguez-Ruano, J. Martinu, F. Vacha, E. Novakova & V. Hypsa (2021) A new symbiotic lineage related to Neisseria and Snodgrassella arises from the dynamic and diverse microbiomes in sucking lice. Molecular Ecology, 30, 2178–2196.

Rihova, J., K. Bell, E. Novakova & V. Hypsa (2022) Lightella neohaematopini: A new lineage of highly reduced endosymbionts coevolving with chipmunk lice of the genus Neohaematopinus. Frontiers in Microbiology, 13:900312. doi: 10.3389/fmicb.2022.900312.

Rihova, J., E. Novakova, F. Husnik & V. Hypsa (2017) Legionella Becoming a Mutualist: Adaptive Processes Shaping the Genome of Symbiont in the Louse Polyplax serrata. Genome Biology and Evolution, 9, 2946–2957.

Rihova, J., R. Vodicka & V. Hypsa (2025) An obligate symbiont of Haematomyzus elephantis with a strongly reduced genome resembles symbiotic bacteria in sucking lice. Applied and Environmental Microbiology, 91(6):e0022025. doi: 10.1128/aem.00220-25.

Ríhová, J., S. Gupta, E. Nováková & V. Hypsa (2024) Fur microbiome as a putative source of symbiotic bacteria in sucking lice. Scientiic Reports, 14(1):22326. doi: 10.1038/s41598-024-73026-2.

Sasaki-Fukatsu, K., R. Koga, N. Nikoh, K. Yoshizawa, S. Kasai, M. Mihara, M. Kobayashi, T. Tomita & T. Fukatsu (2006) Symbiotic bacteria associated with stomach discs of human lice. Applied and Environmental Microbiology, 72, 7349–7352.

Schaub, G. A., A. H. Kollien & C. Balczun. 2012. Lice as Vectors of Bacterial Diseases. In Arthropods as Vectors of Emerging Diseases, ed. H. Mehlhorn, 255–274. Berlin, Heidelberg: Springer Berlin Heidelberg.

Serrato-Salas, J. & M. Gendrin (2023) Involvement of Microbiota in Insect Physiology: Focus on B Vitamins. Mbio, 14(1):e0222522. doi: 10.1128/mbio.02225-22.

Stefka, J. & V. Hypsa (2008) Host specificity and genealogy of the louse Polyplax serrata on field mice, Apodemus species: A case of parasite duplication or colonisation? International Journal For Parasitology, 38, 731–741.

Team, R. C. 2014. R: A language and environment for statistical computing. R Foundation for Statistical Computing, Vienna, Austria.

Trifinopoulos, J., L. Nguyen, A. von Haeseler & B. Minh (2016) W-IQ-TREE: a fast online phylogenetic tool for maximum likelihood analysis. Nucleic Acids Research, 44, W232–W235.

Van der Auwera, G. A., M. O. Carneiro, C. Hartl, R. Poplin, G. Del Angel, A. Levy-Moonshine, T. Jordan, K. Shakir, D. Roazen, J. Thibault, E. Banks, K. V. Garimella, D. Altshuler, S. Gabriel & M. A. DePristo (2013) From FastQ data to high confidence variant calls: the Genome Analysis Toolkit best practices pipeline. Curr Protoc Bioinformatics, 43(1110):11.10.1-11.10.33. doi: 10.1002/0471250953.bi1110s43.

Virtanen, P., R. Gommers, T. Oliphant, M. Haberland, T. Reddy, D. Cournapeau, E. Burovski, P. Peterson, W. Weckesser, J. Bright, S. van der Walt, M. Brett, J. Wilson, K. Millman, N. Mayorov, A. Nelson, E. Jones, R. Kern, E. Larson, C. Carey, I. Polat, Y. Feng, E. Moore, J. VanderPlas, D. Laxalde, J. Perktold, R. Cimrman, I. Henriksen, E. Quintero, C. Harris, A. Archibald, A. Ribeiro, F. Pedregosa, P. van Mulbregt, S. Contributors & S. Contributors (2020) SciPy 1.0: fundamental algorithms for scientific computing in Python. Nature Methods, 17, 261–272.

Wang, W., L. Durden & R. Shao (2020) Rapid host expansion of an introduced parasite, the spiny rat louse Polyplax spinulosa (Psocodea: Phthiraptera: Polyplacidae), among endemic rodents in Australia. Parasites & Vectors, 13(1):83. doi: 10.1186/s13071-020-3957-y.

Yun, J., S. Roh, T. Whon, M. Jung, M. Kim, D. Park, C. Yoon, Y. Nam, Y. Kim, J. Choi, J. Kim, N. Shin, S. Kim, W. Lee & J. Bae (2014) Insect Gut Bacterial Diversity Determined by Environmental Habitat, Diet, Developmental Stage, and Phylogeny of Host. Applied and Environmental Microbiology, 80, 5254–5264.

