## Supplementary figure S1 for "Generalist vs specialist strategy shapes microbiomes in blood feeding parasite *Polyplax serrata*"

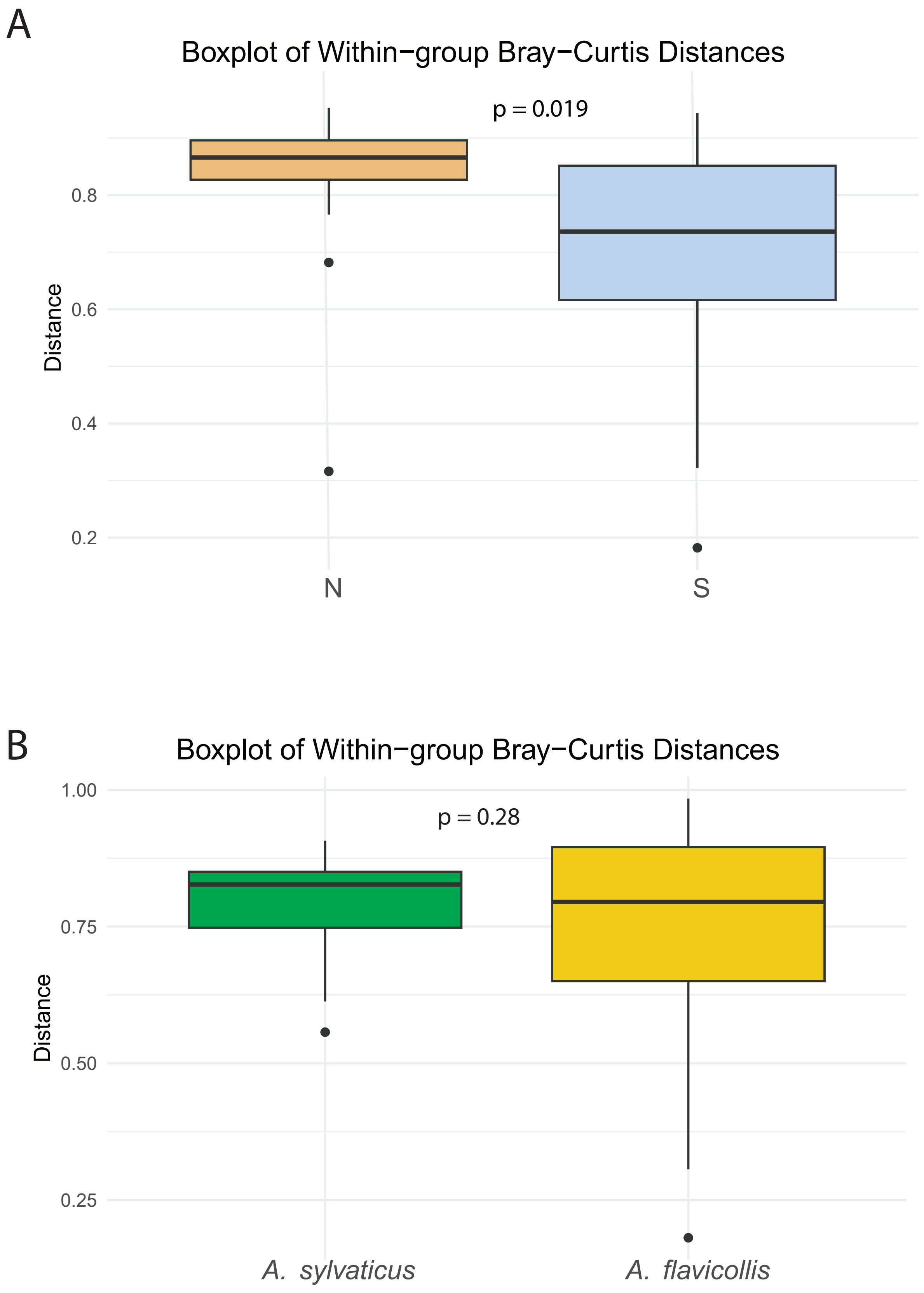


**Supplementary figure S1.** (A) PERMANOVA analysis showing group separation after removal of Bavarian and Saxony samples. (B) PERMANOVA results comparing samples from two host species.
